# Accounting for B cell behaviour and sampling bias yields a superior predictor of anti-PD-L1 response in bladder cancer

**DOI:** 10.1101/2021.03.04.433370

**Authors:** I.A. Dyugay, D.K. Lukyanov, M.A. Turchaninova, A.R. Zaretsky, O. Khalmurzaev, V.B. Matveev, M. Shugay, P.V. Shelyakin, D.M. Chudakov

**Affiliations:** Center of Life Sciences, Skolkovo Institute of Science and Technology, Moscow, Russian Federation.; Genomics of Adaptive Immunity Department, Shemyakin-Ovchinnikov Institute of Bioorganic Chemistry, Russian Academy of Sciences, Moscow, Russian Federation.; Center for Precision Genome Editing and Genetic Technologies for Biomedicine, Institute of Translational Medicine, Pirogov Russian National Research Medical University, Moscow, Russian Federation.; Department of Urology, Federal State Budgetary Institution “N.N. Blokhin National Medical Research Center of Oncology” of the Ministry of Health of the Russian Federation, Moscow, Russia.

## Abstract

Tumor-infiltrating B cells and intratumorally-produced immunoglobulins (IG) play important roles in the tumor microenvironment and response to immunotherapy^1–5^. IgG antibodies produced by intratumoral B cells may drive antibody-dependent cellular cytotoxicity (ADCC) and enhance antigen presentation by dendritic cells^6–8^. Furthermore, B cells are efficient antigen-specific antigen presenters that can essentially modulate the behaviour of helper T cells^9–11^.

Here we investigated the role of intratumoral IG isotype and clonality in bladder cancer. Our results show that the IgG1/IgA ratio offers a strong and independent prognostic indicator for the *Basal squamous* molecular subtype and for the whole ImVigor210 cohort in anti-PD-L1 immunotherapy. Our findings also indicate that effector B cell functions, rather than clonally-produced antibodies, are involved in the antitumor response. High IgG1/IgA ratio was associated with relative abundance of cytotoxic genes and prominence of the IL-21/IL-21R axis suggesting importance of T cell/B cell interaction.

We integrated the B, NK, and T cell components, employing immFocus-like normalization to account for the stochastic nature of tumor tissue sampling. Using a random forest model with nested cross-validation, we developed a tumor RNA-Seq-based predictor of anti-PD-L1 therapy response in muscle-invasive urothelial carcinoma. The resulting PRIMUS (PRedIctive MolecUlar Signature) predictor achieves superior sensitivity compared to PD-L1 expression scores or existing gene signatures, allowing for reliable identification of responders even within the *desert* patient subcohort analyzed as a hold out set.

## Results

### Immunoglobulin isotype composition and clonality

In melanoma, a high IgG1/IgA ratio and large IgG1 clonal expansions—which mainly reflect the presence of clonal IgG1-producing plasma cells in RNA-Seq data—are strongly associated with a positive prognosis^12^. This suggests that cytotoxic tumor-specific antibody production is one of the major mechanisms of melanoma immune surveillance via ADCC and/or antibody-dependent cellular phagocytosis (ADCP)^7,13^. In contrast, for *KRAS*-mutated lung adenocarcinoma, where a high IgG1/IgA ratio is also associated with better prognosis, high clonality of IgG1 repertoire was not associated with long survival^14^. These previous observations suggest the existence of alternative explanations for the association of a better prognosis with a high IgG1/IgA ratio, such as B cell-mediated antigen presentation^9–11^ or direct B cell cytotoxicity^15^.

We investigated bladder cancer (BLCA) patient subcohorts from The Cancer Genome Atlas (TCGA) with distinct mRNA expression-based molecular subtypes in order to identify patients that are more likely to have favorable prognoses while exhibiting a high intratumoral IgG1/IgA ratio (**Fig. 1a, Supplementary Note 1**). This analysis revealed that the *Basal squamous* and *Luminal infiltrated* molecular subtypes show the strongest dependence of patient survival on IgG1/IgA ratio (**Fig. 1b, Supplementary Fig. 1c-f**). *Basal squamous* tumors are generally characterised by high lymphocytic infiltration, including CD8^+^ T cells^16^ and Th1 T cells^17^. Thus, one of the possible explanations for this result could be the association of IgG1 isotype-switching with a type 1 immune response^3,18,19^.

**Figure 1.**
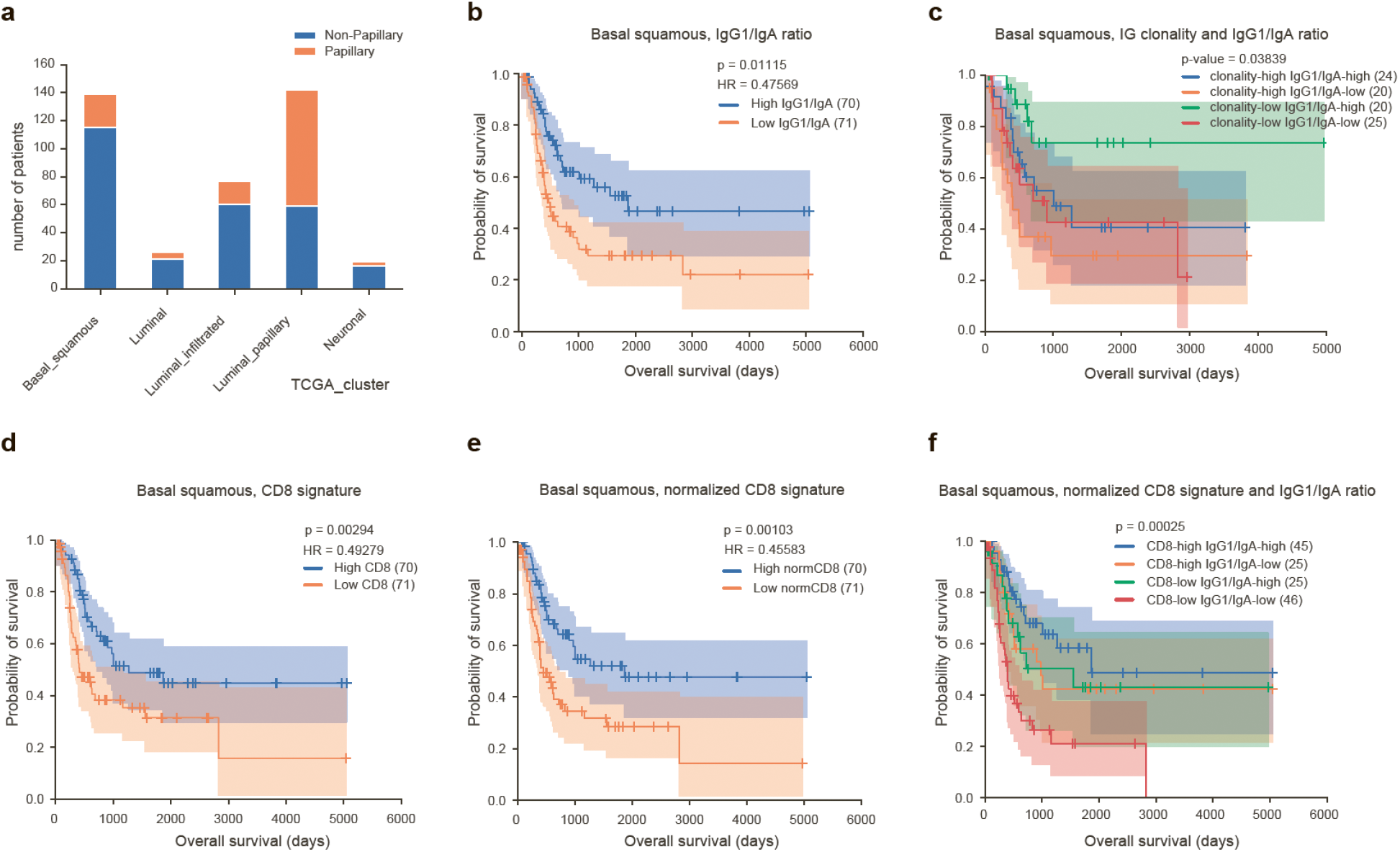
IG repertoire features and normalized CD8 signature predict survival in *Basal squamous* BLCA patients. **a**. Relative overlap of the histological (*Papillary* and *Non-papillary*) and mRNA expression-based molecular subtypes of bladder cancer. **b-f**. Kaplan–Meier overall survival plots for TCGA patients with *Basal squamous* bladder cancer with high and low IgG1/IgA expression ratio (b), a combination of high or low IgG1/IgA expression ratios and high or low IgG clonality (c), high or low CD8 signature (d), high or low CD8 signature calculated after immFocus-like normalization (e), and a combination of high and low IgG1/IgA expression ratios and CD8 signature calculated after immFocus-like normalization (f). Patient cohorts are split by median, with total number of patients shown in parentheses. IgA expression measurements are a sum of IGHA1 and IGHA2.

Next, we studied the clonality of IgG repertoires extracted from TCGA BLCA RNA-seq data using MiXCR^20,21^. We observed that high clonality of the IG repertoire based on RNA-seq data had a non-significant association with a negative prognosis for *Basal squamous* tumours, and was neutral for the full cohort of patients from the TCGA-BLCA dataset (**Supplementary Fig. 2a,b**). However, the combination of IgG1/IgA ratio and IG clonality showed high prognostic value, with the best prognosis associated with high IgG1/IgA and *low* IG clonality (**Fig. 1c**). These results suggest that focused clonal antibody production does not efficiently contribute to immune surveillance of bladder cancer, in contrast to melanoma and similarly to *KRAS*-mutated lung adenocarcinoma. The association of a high IgG1/IgA ratio with a positive prognosis therefore should probably be attributable to other B cell functionalities.

### High IgG1/IgA expression ratio is associated with a cytotoxic immune signature

Next, we aimed to identify the immune processes that are associated with a high IgG1/IgA expression ratio. To this end, we divided *Basal squamous* BLCA patients according to the tertiles based on IgG1/IgA expression ratio, and compared the differential expression of immFocus^22^-normalized genes in patients from high versus low IgG1/IgA tertiles. ImmFocus normalizes the expression of immune-related genes based on the average expression of CD45-correlated genes, allowing one to estimate the relative activity of distinct processes among tumor-infiltrating immune cells independently of the extent of tumor infiltration.

The functions of the normalized genes overexpressed in the high IgG1/IgA subcohort included T cell receptor signaling, CD8^+^ T cell activation, NK cell-mediated cytotoxicity pathways (*CXCL9, CXCL10, CD8A, CD8B, GZMA, GZMB, PRF1, TBX21, IFNG, KLRC2, GNLY*), IL-21-mediated signaling, immune checkpoints (*CTLA4, LAG3, PDCD1*), and B cell receptor signaling, phagocytosis and apoptosis (**Supplementary Fig. 3**; see **Supplementary Table 1** for the full list of positively differentially-expressed genes). This association of a high IgG1/IgA ratio with increased expression of cytotoxic genes suggests a possible correlation between the type 1 T cell response and IgG1 isotype-switching. IL-21 is known to exert an anti-tumor effect by enhancing and supporting CD8^+^ T cell responses^23^, and IL-21 produced by follicular T helper cells (Tfh) promotes B cell proliferation and maturation^24^.

Applying the above technique to the full cohort of TCGA BLCA patients resulted in even more cytotoxic genes being positively associated with a high IgG1/IgA ratio, along with genes involved with CD80, IL-21-mediated signaling, checkpoint regulation, Fc-gamma receptor signaling, receptor-mediated phagocitosis and endocytosis (**Supplementary Fig. 3**).

### Prognostic value of normalized CD8^+^ effector T cell signature and IgG1/IgA ratio

Recently, Mariathasan and colleagues proposed a CD8^+^ effector T cell-associated gene signature that includes *CD8A, CXCL9, CXCL10, GZMA, GZMB, IFNG, PRF1* and *TBX21* genes, which had predictive power in terms of response to anti-PD-L1 immunotherapy^25^ for tumors with an inflamed histological phenotype. When we applied this CD8 signature to the *Basal squamous* TCGA BLCA cohort, we also observed some prognostic value in terms of predicting patient survival (**Fig. 1d**). Notably, after immFocus-like normalization, the same CD8 signature resulted in a much more accurate prognosis (**Fig. 1e**). Moreover, the combination of the IgG1/IgA ratio and normalized CD8 signature had greater prognostic value compared to the CD8 signature alone (**Fig. 1f**). This observation suggests that the IgG1/IgA isotype ratio has independent prognostic value, and is not merely a passive biomarker of type 1 T cell response. Multivariate Cox proportional hazards regression analysis confirmed the independent – and even dominant – contribution of the IgG1/IgA ratio to predicting overall survival of *Basal squamous* BLCA patients (**Supplementary Table 2**).

### Normalized CD8 signature combined with the IgG1/IgA ratio predict survival and response to immunotherapy

We next evaluated the applicability of the normalized CD8 signature and IgG1/IgA ratio to predict the individual response to anti-PD-L1 immunotherapy. We worked with data from the ImVigor210 Phase II clinical trial of atezolizumab in muscle-invasive urothelial carcinoma^25^. The cohort is composed of 345 tissue samples from patients who failed previous platinum-based chemotherapy or previously untreated patients who were ineligible for platinum-based chemotherapy.

The IgG1/IgA ratio alone had moderate predictive value for response to atezolizumab (**Fig. 2a**), and moderate but significant prognostic value (**Fig. 2b**) in terms of overall survival of treated patients. High IG clonality had a non-significant association with a negative prognosis for the full cohort (not shown), but this association was significant for the *Basal squamous* subcohort of treated patients (**Fig. 2c**, subcohort predicted by Kim *et al*.^26^). Low IG clonality combined with a high IgG1/IgA ratio (as in the case with patients from *Basal squamous* TCGA subcohort, **Fig. 1c**) was associated with the best prognosis for the whole ImVigor210 cohort (**Fig. 2d**).

**Figure 2.**
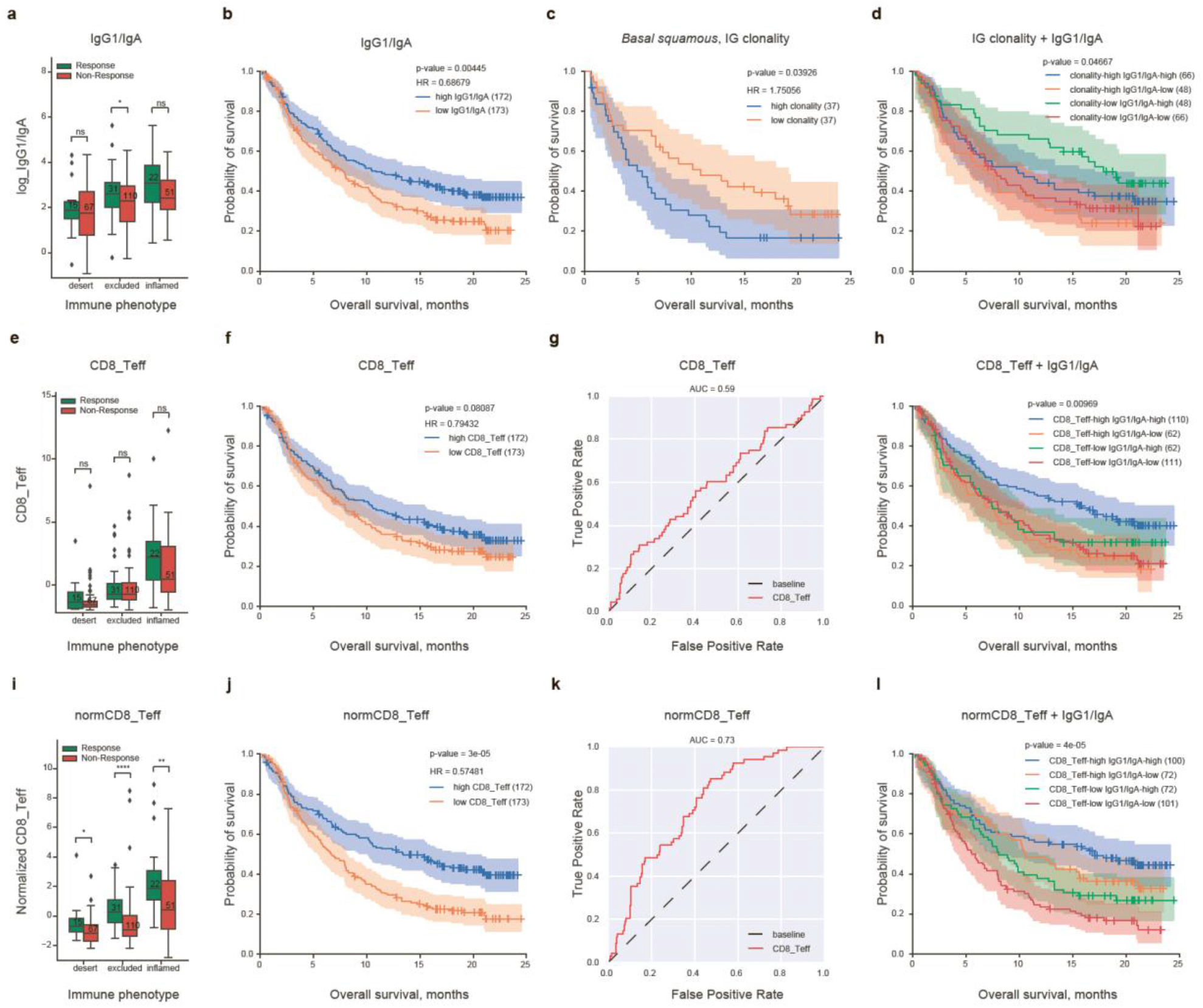
Predictive and prognostic role of different immune features in anti-PD-L1 immunotherapy of bladder cancer, ImVigor210 cohort. **a-d**. Predictive and prognostic role of IgG1/IgA ratio and IG clonality. **e-h**. Predictive and prognostic role of CD8 signature. **i-l**. Predictive and prognostic role of CD8 signature after immFocus normalization. Boxplots (a,e,i) show association with response to anti-PD-L1 immunotherapy for different tumor immune phenotypes. Kaplan–Meier plots show overall survival of patients with different immune features either alone (b,c,f,j) or in combination (d,h,l). Area under receiver operating curve (AUROC) (g,k) shows discriminative ability of a feature to diagnose patients who would benefit from atezolizumab immunotherapy. ns: non-significant; *: 1.00e- 02 < p <= 5.00e-02; **: 1.00e-03 < p <= 1.00e-02 ***: 1.00e-04 < p <= 1.00e-03 ****: p <= 1.00e-04. Except for (c), data are shown for the whole ImVigor210 cohort.

The raw CD8 signature was poorly predictive of response (**Fig. 2e-h**), but immFocus-like normalization dramatically improved its predictive power (**Fig. 2i-k**). The combination of this normalized CD8 signature with IgG1/IgA ratio yielded the best prognostic value (**Fig. 2l**). Multivariate Cox proportional hazards regression analysis again showed a comparable and independent contribution of the IgG1/IgA ratio and normalized CD8 signature in prognosis for ImVigor210 patients (**Supplementary Table 2**).

To elucidate which immune-related features are associated with response to immunotherapy after immFocus normalization, we split ImVigor210 patients into two equally-sized groups while maintaining the proportion of responders and non-responders. One group was used to search for genes that were differentially expressed in responders. We performed gene set enrichment analysis (GSEA) of the top 100 differentially-expressed genes. Among the immune-related features associated with response, we found a Ras-independent pathway in NK cell-mediated cytotoxicity including *KLRC2, KLRC3* and *KLRC4*. We confirmed these results in the second group of patients. Notably, *KLRC1* and *KLRC2* were also positively differentially expressed in TCGA BLCA patients with a high IgG1/IgA isotype ratio (**Supplementary Fig. 3** and **Supplementary Table 1**).

### Integrative predictive modeling of response to anti-PD-L1 immunotherapy

Immunohistochemical measurement of PD-L1 expression in tumor samples is currently the only approach that can identify patients who may have a higher chance of responding to immunotherapy. There are various metrics that account for the presence of PD-L1 on the surface of immune cells (ICs) and tumor cells. The SP142 Ventana immunohistochemistry assay was used in the ImVigor210 trial. In this assay, antibodies to the PD-L1 C-terminus are used, and the scoring system (ICA) is calculated as the percentage of PD-L1-positive ICs in a given tumor area^27,28^. Currently the cut-off for first-line therapy is 5%, but the predictive value is relatively low (see **Fig. 3a-c** and Refs 29–31).

**Figure 3.**
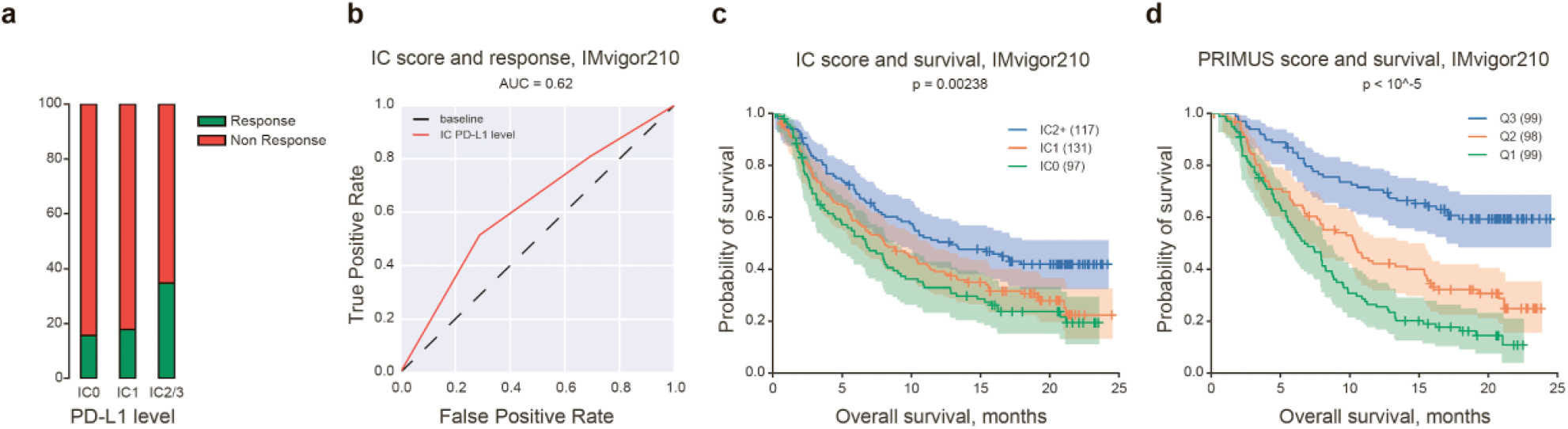
Prognostic value of immunohistochemical PD-L1 measurements among tumor-infiltrating immune cells, ImVigor210 cohort. **a**. Percentage of responders and non-responders among patients divided based on the amount of PD-L1^+^ ICs (IC0: <1%, IC2: 1-5%, IC2/3: >5%) as assessed by the SP142 immunohistochemistry assay. **b**. Receiver operating curve (ROC) representing the discriminative ability of PD-L1^+^ IC counts in biopsies to identify patients who would benefit from atezolizumab immunotherapy. **c**. Kaplan-Meier survival plots for patients with different counts of PD-L1^+^ ICs. **d**. Kaplan-Meier survival plots for patients divided according to predicted probability of response by PRIMUS. Patients are split by tertiles. Note that although the model was trained using different regularization terms and is thus considered to be protected against overfitting, this panel formally includes the data the model was trained on and is therefore shown for illustrative purposes only.

To develop a stronger gene expression-based predictor for rational patient stratification, we used a random forest model, with performance evaluated using a nested cross-validation approach^32^ (**Fig. 4**).

**Figure 4.**
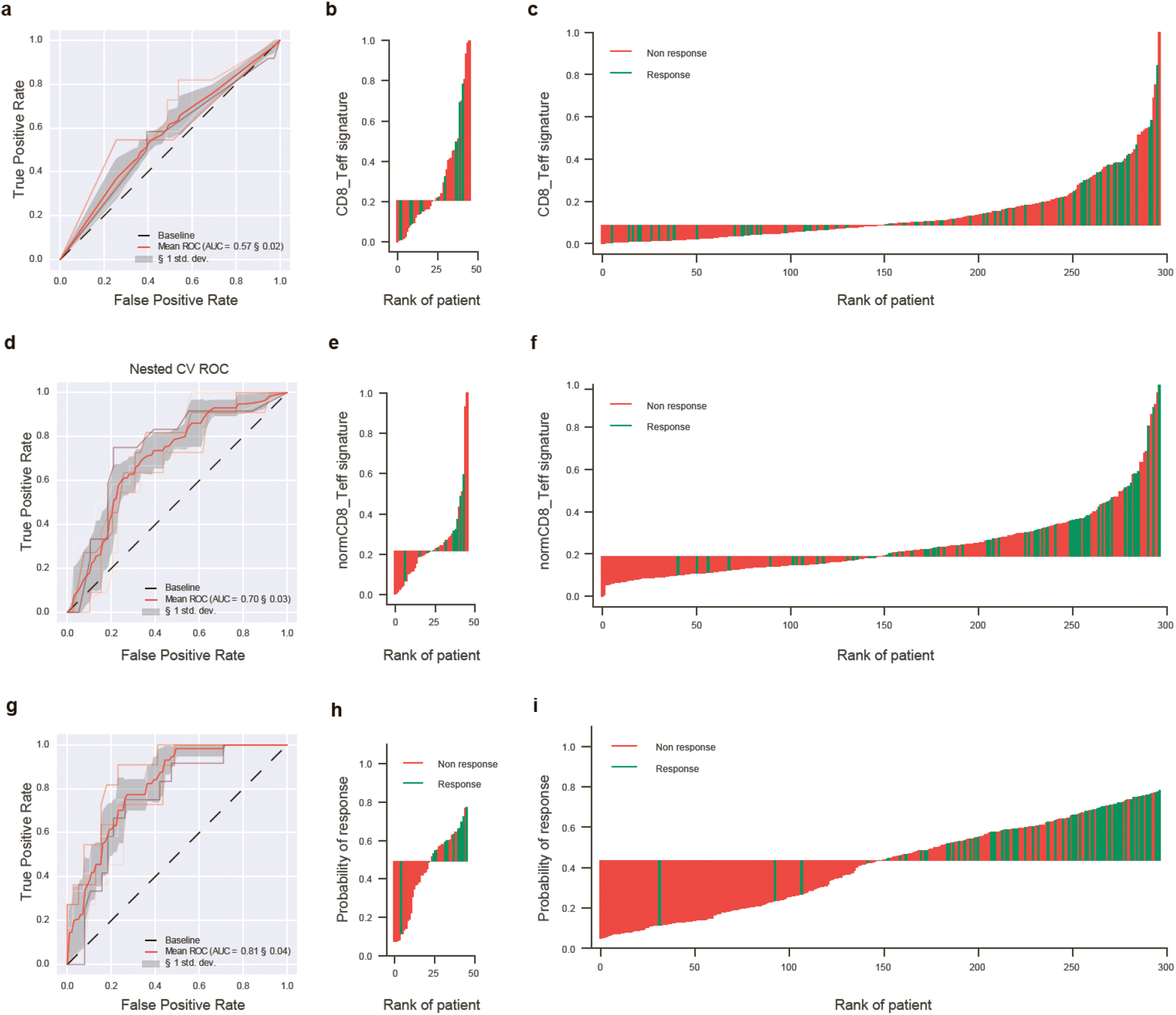
Predictive value of CD8 signature, immFocus-normalized CD8 signature, and integrative random forest model. **a-c**. CD8 signature (*CD8A, CXCL10, CXCL9, GZMA, GZMB, IFNG, PRF1, TBX21*). **d-f**. immFocus-normalized CD8 signature. **g-i**. Model based on refined immFocus-normalized CD8 signature combined with immFocus-normalized KLRC-NK signature, immFocus-normalized IL-21, and GNLY expression. AUROC (a,d,g) shows the discriminative ability of a signature to identify patients who would benefit from atezolizumab immunotherapy. Waterfall charts show distributions of responders and non-responders according to corresponding ranked-feature values for holdout sets (b,e,h) and the whole patient cohort (c,f,i). Although the model was trained using different regularization terms and is considered to be protected against overfitting, panel (i) formally includes the data the model was trained on and is therefore shown for illustrative purposes only.

First, to set a baseline for our model’s performance, the model was trained using solely unnormalized CD8 signature. We got an area under the receiver operating curve (AUROC) similar to one made from the ranked values of the signature (compare **Fig. 2g** and **Fig. 4a**), which doesn’t differ much from the random algorithm. The mean nested cross-validation F1 score, which reflects the overall balance between precision and recall (where closer to 1 is better), was 0.344 ± 0.06.

Next we trained the model using the immFocus-normalized CD8 signature. We got an AUROC that was much greater than one we got for the unnormalized signature, and similar to the AUROC from the normalized signature itself (compare **Fig. 2k** and **Fig. 4d**). The mean nested cross-validation F1 score was 0.465 ± 0.05.

Before selecting the final set of input features, we further refined the CD8 signature by fitting the model on each gene from the signature separately rather than in combination. The four genes with highest importance were selected: *CXCL9, CXCL10, CD8A, GZMB*. To appropriately estimate feature importance, we also analysed them for the presence of multi-collinearity in the data. For each variable, we calculated a variance inflation factor that was <5, indicating that we had no multi-collinear input parameters. We also excluded from the ultimate model several parameters that demonstrated no significant predictive value, including IG clonality, *IL21R* and *CD80*.

Finally, we integrated features including immFocus-normalized CD8 signature (*CXCL9, CXCL10, CD8A, GZMB*), immFocus-normalized KLRC-NK signature (*KLRC2, KLRC3, KLRC4*), immFocus-normalized *IL21* and *GNLY* expression, IgG1/IgA isotype ratio, and non-normalized *TGFB1* expression^25^.

The performance of the final PRIMUS (PRedIcitive MolecUlar Signature) model was higher than that of the model trained on the immFocus-normalized CD8 signature (**Fig. 4g-i**). The mean nested cross-validation F1 score was 0.495 ± 0.05.

To confirm the validity of the model, we compared the performance of PRIMUS with the support vector machine (SVM)-based model with a linear kernel. This is a simple model that is well-designed for discriminating linearly-separable data, and is unlikely to overfit complex data. The results were comparable with those obtained with PRIMUS, with a mean nested cross-validation F1 score of 0.45 ± 0.05. This indicates that our final model was not overfitted.

### Feature importance and interaction analysis

We next retrained PRIMUS on the full set of ImVigor210 data to explore interactions between and the importance of input features using SHAP^33,34^, a game theory approach. First, we compared the importance of our input variables to randomly generated numbers. Each of the PRIMUS variables was shown to be more important than randomly-generated numbers, but normalized CD8 signature had the most importance (**Fig. 5a,b**).

**Figure 5.**
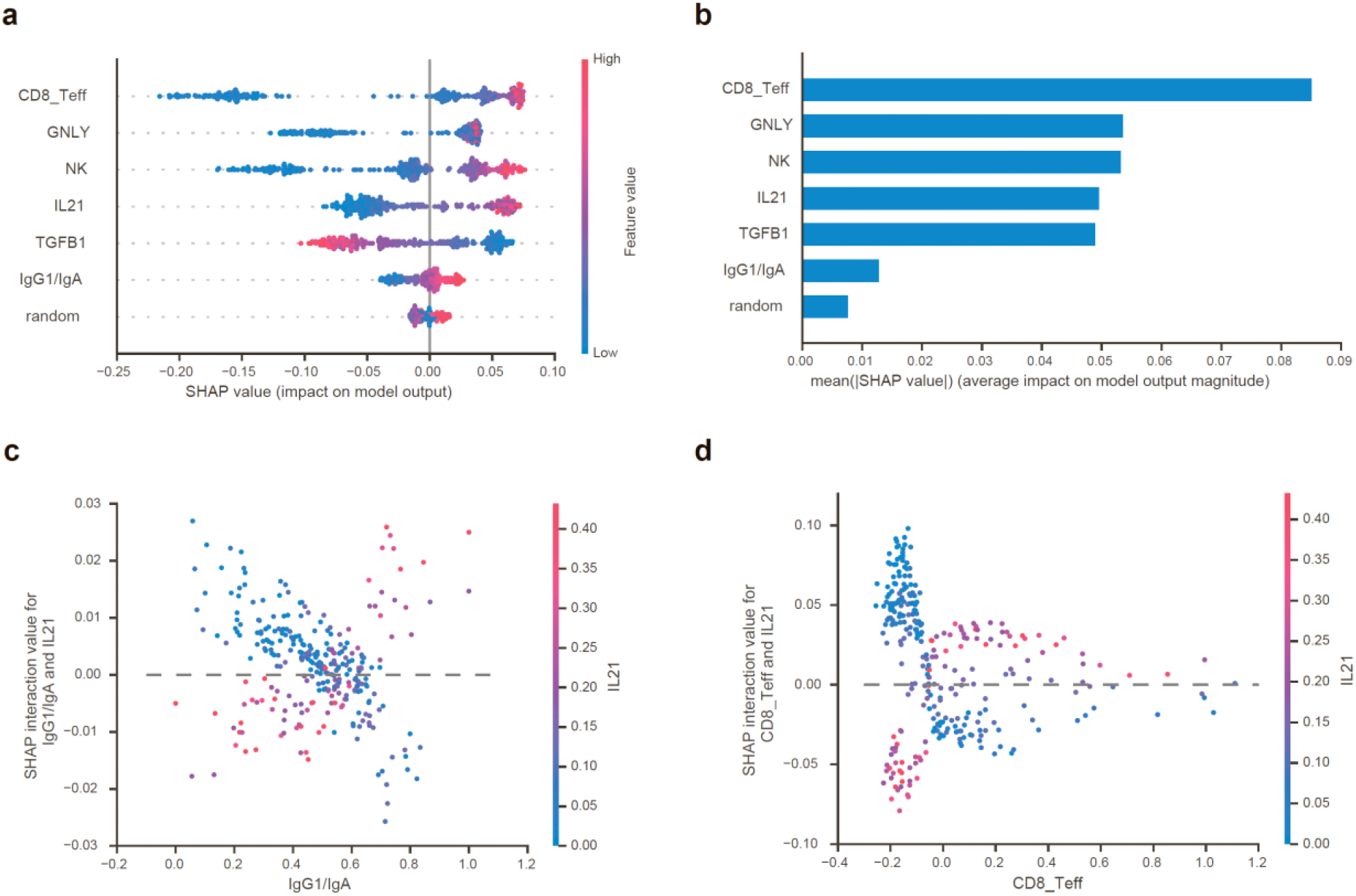
PRIMUS feature importance and interactions. **a, b**. PRIMUS feature importance compared with randomly-generated numbers **c**. Impact of interaction between IgG1/IgA ratio and *IL21* expression. **d**. Impact of interaction between CD8 signature and *IL21* expression.

We also identified interesting interactions between the variables. In particular, a high IgG1/IgA ratio was more valuable for response prediction when combined with high *IL21* expression (**Fig. 5c**), supporting the involvement of T and B cell interaction in response. There were also interactions between CD8 signature and *IL21* expression, with high expression of *IL21* becoming more of a determining factor with higher values of the CD8 signature (**Fig 5d**). We also observed that only extremely high values of IgG1/IgA ratio are important for response prediction, in contrast with CD8 signatures, for which we can see strong discriminative power (**Fig 5a, Supplementary Fig. 4**).

### Integrative predictor demonstrates high efficiency on the “desert” cohort

Three basic tumor immune phenotypes can be distinguished based on biopsy histology. *Inflamed* phenotype is characterized by the abundant infiltration of tumor parenchyma with CD4^+^ and CD8^+^ T cells, often accompanied by other immune cells, including immunosupressive subtypes. *Excluded* phenotype is characterised by the localization of multitudes of immune cells in the tumor stroma instead of the parenchyma, surrounding nests of tumor cells. Finally, the *desert* phenotype is characterized by a non-inflamed tumor microenvironment, with few or no T cells in either the parenchyma or stroma of the tumor^35,36^. It is particularly hard to predict response among desert patients^37^. In particular, the CD8 signature, TGFβ response signature, and tumor mutational burden all failed to predict response in the *desert* patients in ImVigor210 study^25^.

Notably, PRIMUS predicted response in *desert* patients of the ImVigor210 trial after results being extrapolated to a whole cohort (**Fig. 6a**), with the caveat that although our model was generally protected against overfitting, this analysis formally included the data that the model was trained on.

**Figure 6.**
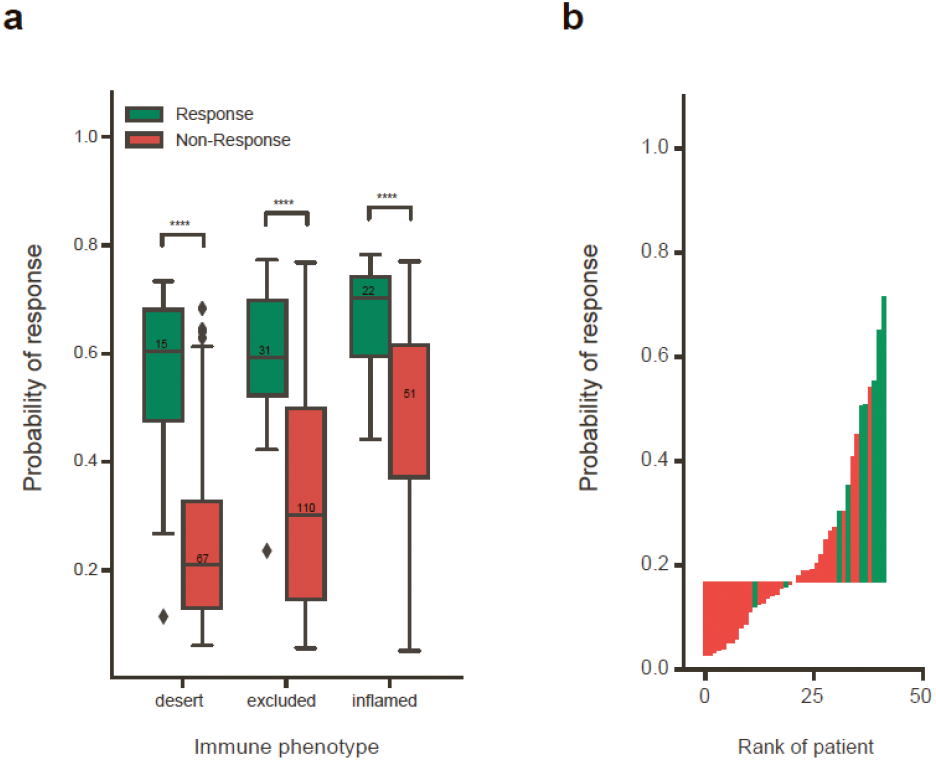
Performance of PRIMUS on patients with “desert” immune phenotype. **a**. Boxplots show probability of response to anti-PD-L1 immunotherapy for different tumor histological immune phenotypes. ***: 1.00e-04 < p <= 1.00e-03 ****: p <= 1.00e-04. Note that the plotted data include those the model was trained on. **b**. “Desert” hold-out set, 41 patients, split by median.

To verify if our model can efficiently predict response among desert tumor phenotypes in a completely blind mode, we trained PRIMUS on the ImVigor210 cohort, leaving 41 *desert* patients as a hold-out set. Remarkably, PRIMUS successfully predicted response for these patients (**Fig. 6b**), thus solving one of the most difficult puzzles in predicting the response to anti-PD-L1/PD1 immunotherapy.

## Discussion

There are several efforts underway to develop rational prognostic and predictive signatures for BLCA patients based on the analysis of differentially expressed genes^38–41^or immunohistochemistry-based classification systems^42^. However, most of these do not take into account the stochastic aspects of tumor sampling. Any tumor sample from a given patient will most likely not fully represent the entirety of a highly heterogeneous tumor tissue^43,44^. From this point of view, the accuracy of an immune gene signature calculation is intrinsically limited by the stochastic nature of tissue sampling, where the observed variability in immune gene expression levels results from the amount of immune cells that happen to infiltrate a particular tissue fragment being sampled. Furthermore, there are few cases of studying specific biological processes that might be involved in survival and immunotherapy response for BLCA patients, that could help in developing the rational prediction algorithms^45^.

There is a solid rationale for using CD8^+^ T cell-specific gene expression^46^ and immunohistochemical^47^ features to predict treatment response. However, there is also no doubt that an effective predictor of response should take into account diverse components of the immune system, and that such predictors may differ greatly for different types of cancer^48^. In particular, multiple studies have demonstrated substantial involvement of CD4^+^ T cells^49,50^, NK cells^51^, and B cells^1–5^ in cancer immunosurveillance and immunotherapy response^1,2^.

In our approach, we combined these parameters of tumor microenvironment, and accounted for the relative representation of each of the immune processes. We hypothesized that the relative activity of distinct immune components among the tumor-infiltrating immune cells may prevail over the infiltration level *per se* in terms of prognostic and predictive value in the context of immunotherapy. We reasoned that the estimation of such relative activity of immune processes based on normalization against pan-leukocyte genes would essentially neutralize the artificial variability in expression levels resulting from random tissue sampling. Notably, the resulting normalized CD8 signature produced a major advance in terms of predicting treatment response and survival after immunotherapy (**Fig. 2i-k, Fig. 4d-f**). The IgG1/IgA ratio, which essentially reflects B cell behavior (**Supplementary Fig. 3**), contributed independently and prominently improved the prognostic value (**Fig. 2l**).

By combining rationally pre-selected parameters, including normalized CD8 signature, IgG1/IgA ratio, and a limited number of genes involved in NK cell response and T cell/B cell interaction, we were able to develop PRIMUS: a model that efficiently predicts response to anti-PD-L1 immunotherapy in muscle-invasive urothelial carcinoma (**Fig. 4g-i**), including tumors with the especially challenging *desert* phenotype (**Fig. 5**). Pending further validation, we hope to pursue clinical implementation of our approach and its derivatives in the near future.

Furthermore, our results show considerable future potential for predicting the response to immunotherapy using transcriptomic data. By building on a deeper understanding of the immune processes underlying an effective antitumor response and using relevant statistical approaches, we can make further progress in developing predictors of response to a given therapy or combinations thereof.

## Methods

### Data used

Patient data from TCGA BLCA include 412 cases, for 408 of which RNA-seq data are available^52^. Cases contain data on both tumors and healthy tissues. For some patients, multiple replicates of tumor samples are present. We downloaded FPKM-UQ files from the GDC data portal. These represented 433 tumor samples, including replicates; we used only one replicate for each patient, randomly selected. The data was then transformed to TPM. We also used raw RNA-seq reads from TCGA-BLCA cohorts to extract B cell receptor sequences. Patient data from the ImVigor210 clinical trial include 345 RNA-seq tissue transcriptomes from patients who failed previous platinum-based chemotherapy or previously untreated patients who were not eligible for platinum-based chemotherapy. The data was downloaded from EMBL-EBI^53^. Abundances of transcripts from bulk RNA-Seq data were quantified using Kallisto^54^.

### ImmFocus normalization

ImmFocus normalization was proposed by Teltsh *et al*.^22^ with modifications. We first selected an immune normalized gene set (INGS): the group of genes for which the Spearman correlation coefficient with *PTPRC* (CD45) was > 0.9. Next, the sample-specific normalization factor (f_INGS_) was calculated for each sample as the averaged expression of genes from INGS, and the first normalization was performed. The normalization coefficient for genes included in INGS did not count itself to avoid self-normalization, and was calculated as the averaged expression of the remaining genes. Then we selected genes from INGS for which the ratio between the coefficient of variation before and after the first normalization was < 0.8 and used those genes as the final INGS. The second normalization was performed using the final INGS.

### Differential expression analysis

Differential expression analysis was performed using the Mann-Whitney U test to find differentially-expressed genes in two samples of patients. Obtained p-values were adjusted with the Benjamini-Hochberg procedure. The fold change was calculated for each gene as the ratio of the median expression in the two samples.

### Statistical analysis

Survival analysis was performed with the lifelines^55^ Python library. Survival plots were created using the Kaplan-Meier estimator. Cox proportional hazard models were fitted on either features or features with interaction value. The relative reliability of models was estimated by the Akaike information criterion and concordance index. The Cox AUROC was calculated with the Python scikit-learn library. For multiple comparisons, correction was performed using the Benjamini-Hochberg procedure^56^. Group comparison in boxplots was performed with the Kruskal-Wallis test. All statistical calculations were performed using Python. P values < 0.05 were considered statistically significant.

### Clonality analysis

We obtained IGH CDR3 repertoires from raw RNA-Seq data using MiXCR 3.0^20,21^ in RNA-seq mode. Data was pre-filtered based on 15-mer nucleotide matches to V/J segments of immune receptors to speed up search. Samples with less than 300 IGH CDR3-related reads were omitted from the analysis. IG CDR3 repertoires were downsampled to 300 randomly-chosen reads for normalization purposes. IG clonality was calculated as [1 - normalized Shannon-Wiener index]^58^ using VDJtools^57^ software.

### Gene signatures

Gene signatures were calculated as the first principal component of principal component analysis (PCA) performed with z-score transformed expressions of input genes. The calculations were performed with PCA from the Python scikit-learn library.

### Random forest model

The random forest was performed with the RandomForestClassifier from the Python scikit-learn library. During training, we selected 15% of the initial data as a holdout set, preserving the proportions of response or non-response to immunotherapy. Hyperparameters were tuned with RandomizedSearchCV and GridSearchCV from the Python scikit-learn library using five-fold cross-validation. The number of estimators in the model was 50. The model was trained using F1 score as a measure of quality: F1=2*precision*recall/(precision+recall). The model performance was evaluated with five-inner/five-outer nested cross-validation. This approach, unlike regular cross-validation, assumes to fit the model using two nested loops, where the inner one is used for hyperparameter optimization and model selection, as with regular cross-validation, while the outer one is used to split the data into training and test folds in order to estimate the performance criterion. We decided to use nested cross-validation because of the small size of the available data points, unbiased generalization performance estimation, and prevention of selection bias.

## Funding

Supported by grant of the Ministry of Science and Higher Education of the Russian Federation º 075-15-2020-807.

## Acknowledgements

We are grateful to Michael Eisenstein for his valuable help in editing the manuscript.

## Conflict of interest

Authors declare no competing financial interests.

## Supplementary Note 1. BLCA subtypes

Bladder cancer is divided into three disease states: non-muscle invasive, muscle-invasive, and metastatic. There are also several variants within these categories, both in terms of histology and gene expression, with different clinical significance^59^.

In terms of histological subtypes, ~90% of bladder cancers are conventional urothelial (transitional cell) carcinomas (UC) and its histologic variants, with the remainder characterized as non-urothelial carcinoma, including squamous and glandular neoplasms. UC can be further subdivided into papillary and non-papillary carcinomas. Non-papillary carcinomas can exhibit variable spectrum of histological features, including squamous differentiation^59,60^.

In this work, we used TCGA 2017 mRNA clusters^16^ that include *Luminal infiltrated, Luminal papillary, Luminal, Basal squamous* and *Neuronal* subtypes. The *Luminal papillary* subtype is characterized by enrichment in tumors with papillary morphology as well as a high rate of *FGFR3* mutations. The *Luminal infiltrated* subtype can be distinguished from other luminal subtypes by increased lymphocytic infiltration and increased expression of PD-L1 and PD-1. The *Luminal* subtype has the highest expression levels of uroplakins and genes that are expressed by terminally-differentiated urothelial umbrella cells. The *Basal squamous* subtype is characterised by high expression of basal and stem-like markers and shows the strongest immune gene expression signature, including T cell markers and pro-inflammatory genes. The *Neuronal* subtype shows relatively high expression of neuronal differentiation genes as well as neuroendocrine markers.

**Supplementary Figure 1.**
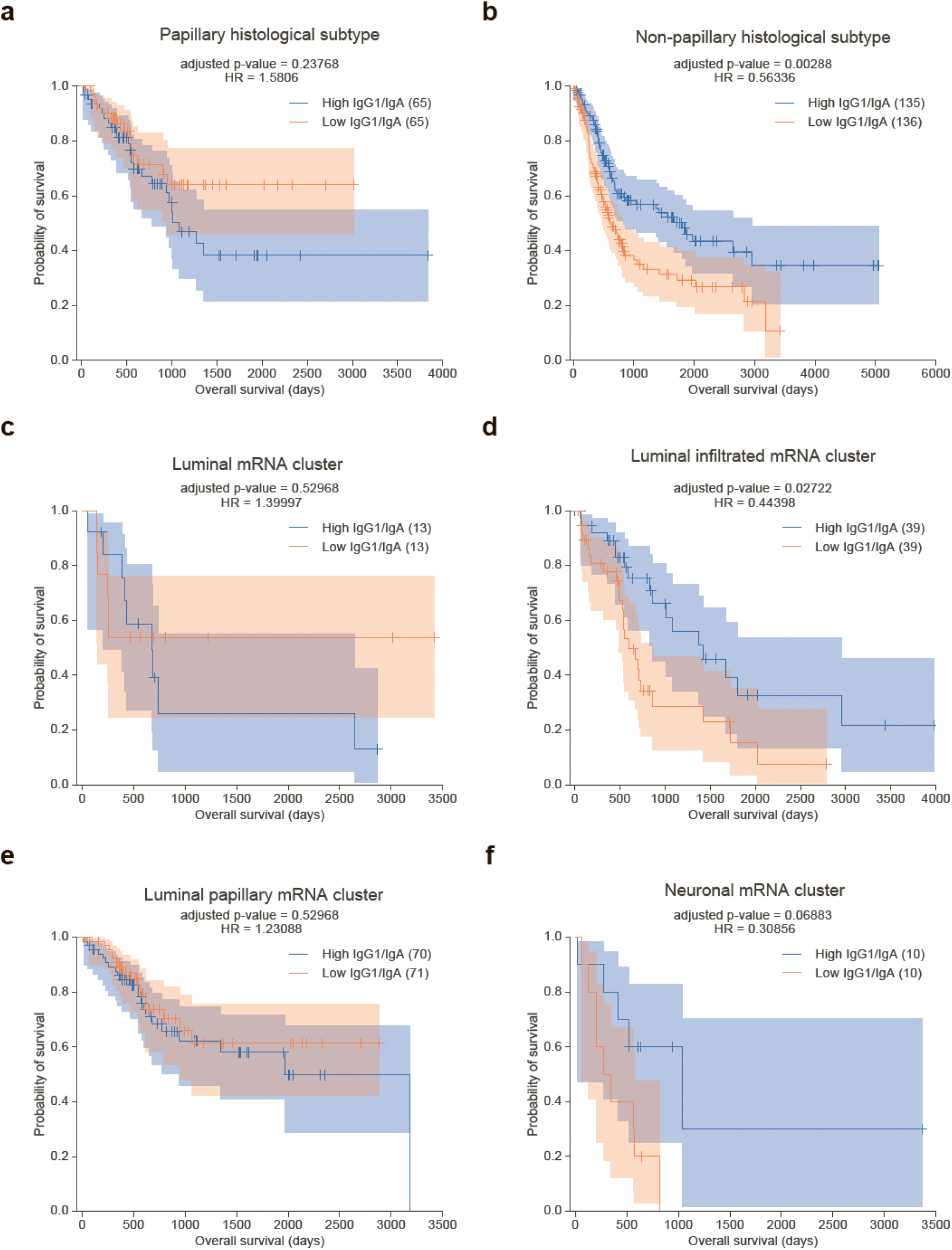
Role of IgG1/IgA expression ratio in bladder cancer subtypes. Kaplan–Meier overall survival plots for patients with **a**. *Papillary*, **b**. *Non-papillary*, **c**. *Luminal*, **d**. *Luminal infiltrated*, **e**. *Luminal papillary*, or **f**. *Neuronal* tumor subtypes as a function of IgG1/IgA expression ratio, where IgA is a sum of IGHA1 and IGHA2. Patient cohorts are split by median, with total numbers in parentheses.

**Supplementary Fig. 2.**
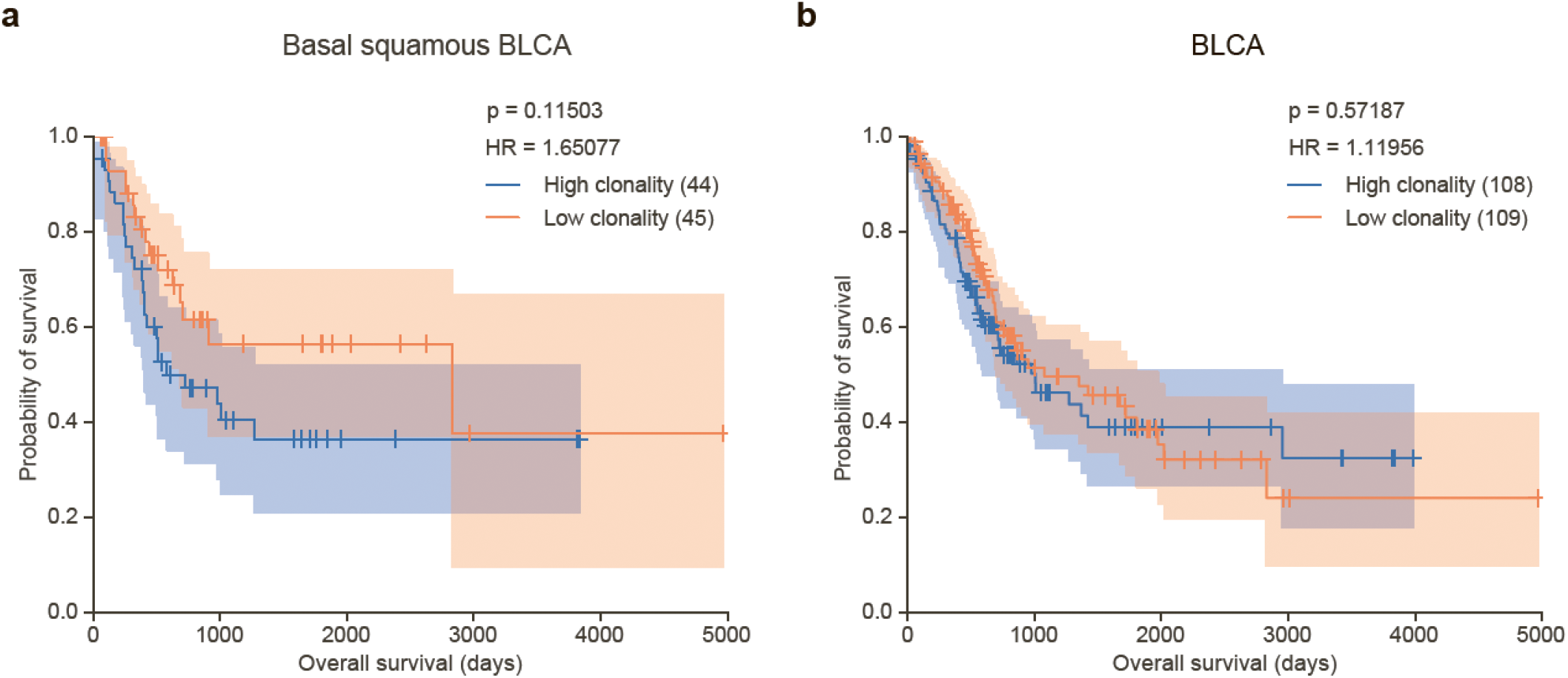
IGH clonality and survival in TCGA BLCA patients. **a, b**. Kaplan–Meier overall survival plots for TCGA *Basal squamous* BLCA patients (a) and the full cohort from the BLCA dataset (b) with high and low IGH clonality. Patient cohorts are split by median, with total numbers in parentheses.

**Supplementary Figure 3.**
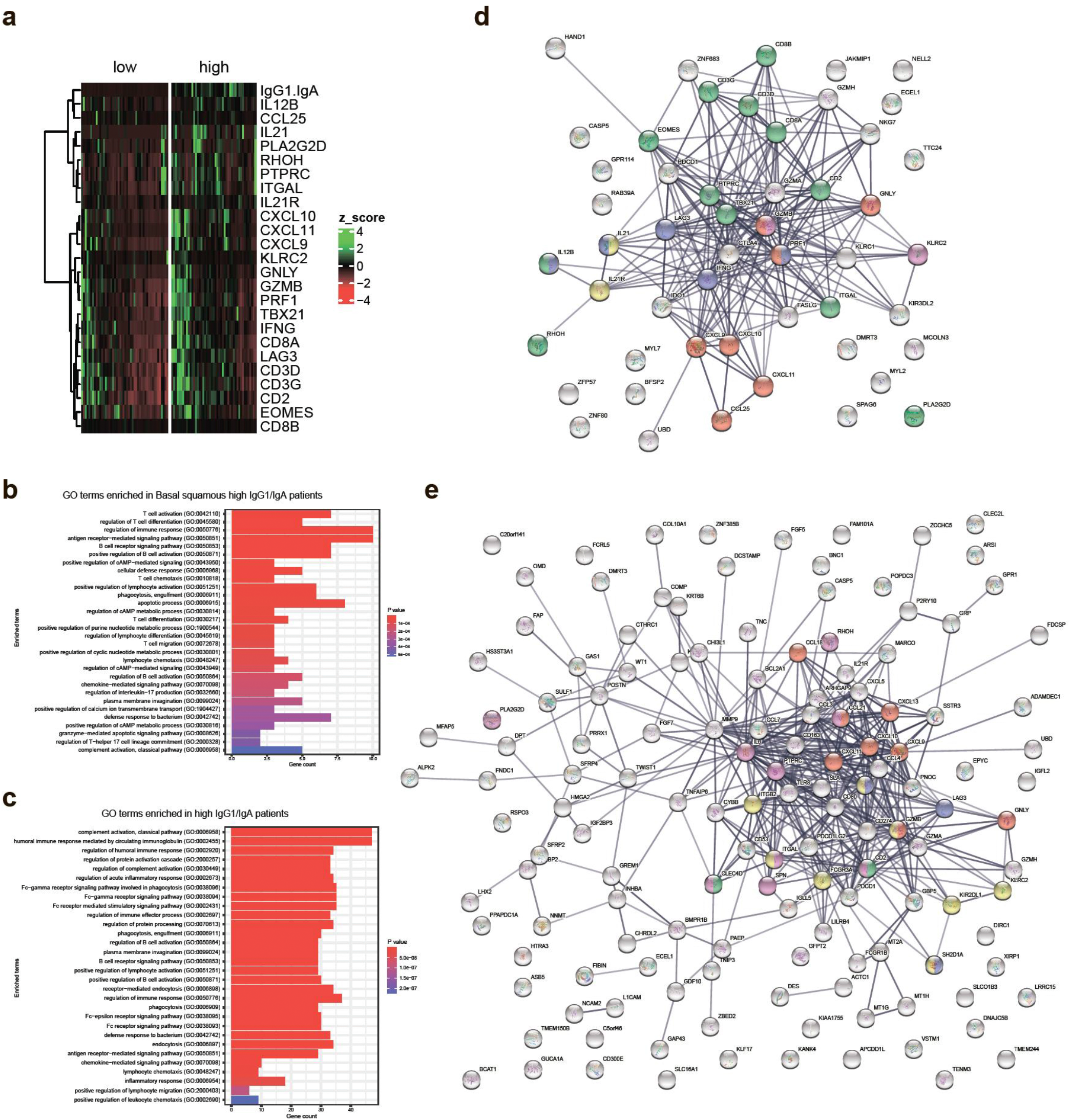
Differential gene expression associated with high IgG1/IgA ratio. **a**. Heatmap of expression of immFocus-normalized (z-scored) immune genes in the sub-cohort of *Basal squamous* patients. Columns are grouped by low or high IgG1/IgA ratio. **b,c**. Gene Ontology (GO) immune processes represented among the normalized genes that are overexpressed in IgG1/IgA-high *Basal squamous* BLCA patients (b) and IgG1/IgA-high subcohort of all TCGA BLCA patients (c). **d**. GO immune processes overexpressed in IgG1/IgA-high *Basal squamous* TCGA BLCA patients visualized with STRING^61^. Red - cell killing, blue - positive regulation of cell killing, green - T cell activation, yellow - IL21-mediated signaling pathway, purple - NK cell mediated immunity. **e**. GO immune processes overexpressed in IgG1/IgA-high subcohort of all TCGA BLCA patients visualized with STRING. Red - cell killing, blue - positive regulation of cell killing, purple - T cell activation, green - positive regulation of myeloid dendritic cell activation, yellow - NK cell mediated cytotoxicity.

**Supplementary Figure 4.**
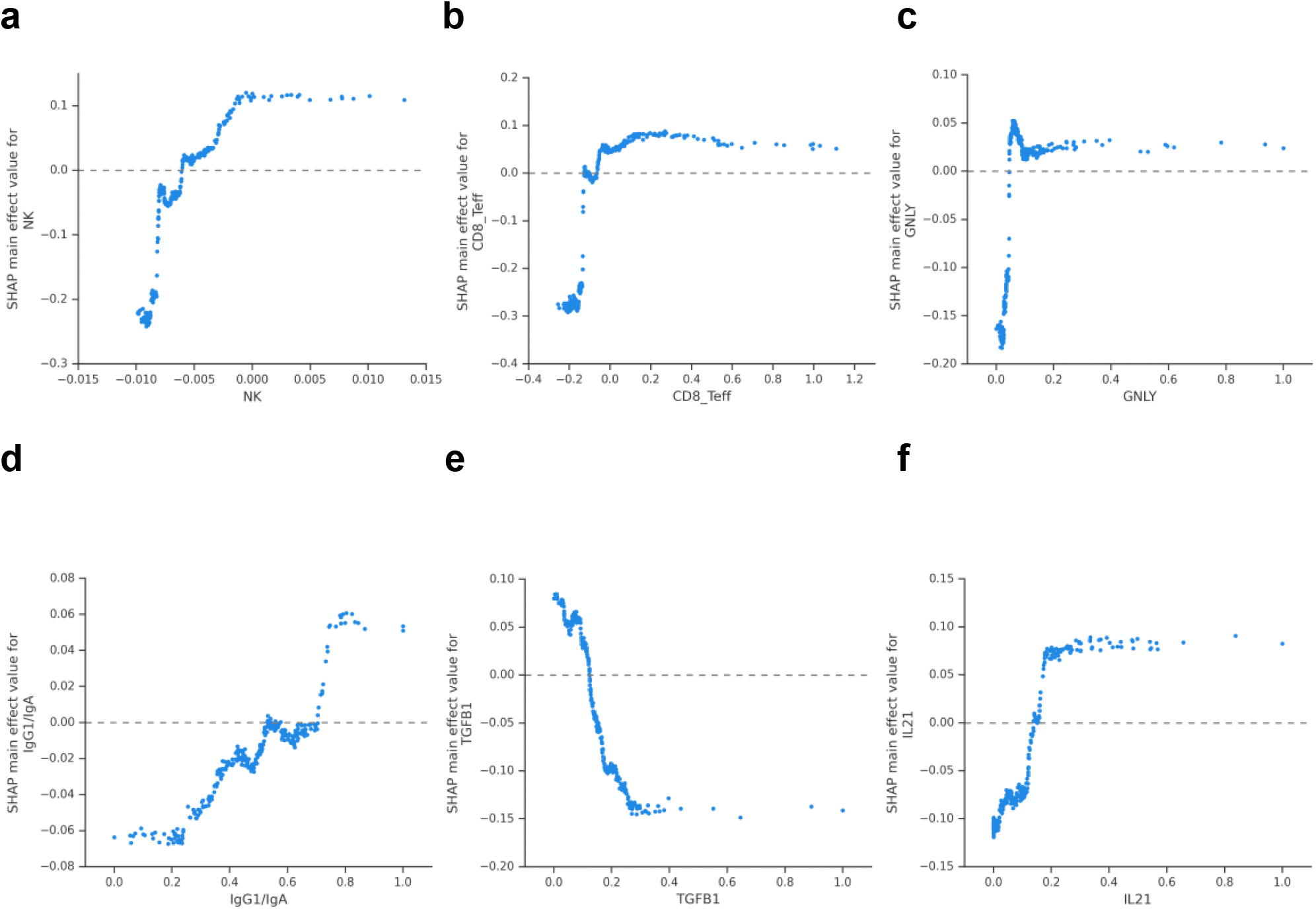
Individual effects of PRIMUS features on response prediction. Individual effect for **a**. NK signature, **b**. CD8 signature, **c**. *GNLY* expression, **d**. IgG1/IgA ratio, **e**. *TGFB1* expression, and **f**. *IL21* expression.

## Notes

### Competing Interest Statement

The authors have declared no competing interest.

